# FastPG: Fast clustering of millions of single cells

**DOI:** 10.1101/2020.06.19.159749

**Authors:** Tom Bodenheimer, Mahantesh Halappanavar, Stuart Jefferys, Ryan Gibson, Siyao Liu, Peter J. Mucha, Natalie Stanley, Joel S. Parker, Sara R. Selitsky

## Abstract

Current single-cell experiments can produce datasets with millions of cells. Unsupervised clustering can be used to identify cell populations in single-cell analysis but often leads to interminable computation time at this scale. This problem has previously been mitigated by subsampling cells, which greatly reduces accuracy. We built on the graph-based algorithm PhenoGraph and developed FastPG which has the same cell assignment accuracy but is on average 27x faster in our tests. FastPG also has higher cell assignment accuracy than two other fast clustering methods, FlowSOM and PARC.

**Availability:** FastPG is available here: https://github.com/sararselitsky/FastPG

---

Mass cytometry measures proteins abundances at the single-cell level. This technology can measure 100,000-200,000 cells per sample and up to around 40 protein markers simultaneously^1^. Unsupervised clustering is a common task in single-cell data analysis, with the goal of unbiasedly identifying known and unknown cell-types based on protein markers. Clustering cells across multiple samples can be optimal for some experiments to gain a deeper understanding of how a particular cell population changes in a disease state. Most clustering algorithms cannot efficiently handle a very large number of collected cells. These algorithms are severely restricted by computational time and available system memory. Common approaches to mitigate these issues include subsampling and meta-clustering^2-4^. The disadvantage to both is loss of information and lower potential to discover rare cell types^4^.

A commonly used and robust method for mass cytometry data is PhenoGraph^5^, which uses a graph-based approach to unbiasedly identify clusters, or cell types. Here we present FastPG (Figure 1A), a modified version of PhenoGraph which was developed to maximize computational efficiency with no loss of cell assignment accuracy. PhenoGraph is composed of three main steps: (1) creating a k-nearest neighbor network (kNN) of single cells using a distance measurement calculated from their protein marker abundances, (2) adding weights to the network through calculating Jaccard index, and (3) partitioning cells into coherent cell populations using the Louvain algorithm^6^. We modified PhenoGraph as follows: first, we replaced the kNN step with a fast kNN approximation, Hierarchical Navigable Small World (HNSW), which uses logarithmic scaling due to the hierarchical structure of the search space^7^. We next parallelized the Jaccard index step for multithreaded execution, and lastly, we replaced the Louvain algorithm with a fast parallelized version, Grappolo^8^.

**Figure 1.**
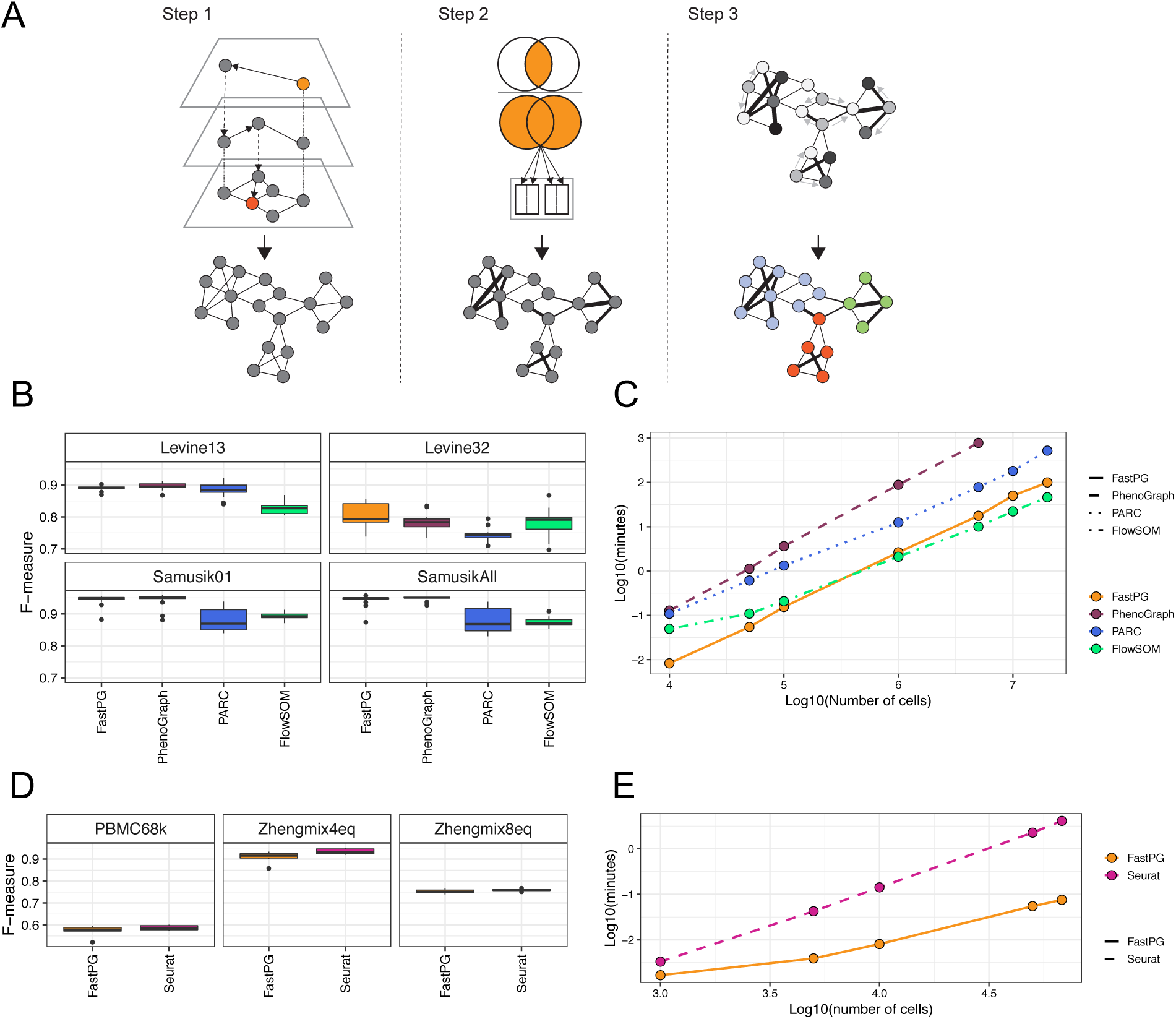
**(A)** Schematic which depicts FastPG. Top row shows a representation of FastPG’s modifications. Bottom row shows the output of each step. **Step 1.** kNN was replaced with the kNN-approximation HNSW, which has logarithmic scaling due to the hierarchical structure of the search space (depicted). The output of this step is a network of cells, where each node is a cell and neighbor are connected by an edge. **Step 2.** Modification of the Jaccard index step to run in parallel, depicted as being distributed to each thread of the CPU. This step adds weights to the network, which are represented as different edge thicknesses. **Step 3.** Louvain algorithm was replaced with a parallelized version, Grappolo, which implements several heuristics for efficient parallelization for fast community detection. The graph coloring heuristic is depicted, where different shades of gray represent different colors assigned to nodes (no two neighbors should receive the same color). The output of this step is the assignment of cells to communities, which was depicted with different colored nodes. **(B)** Boxplot displaying the F-measure for four mass cytometry “gold standard” datasets. **(C)** Runtime comparisons between PhenoGraph, FastPG, PARC, and FlowSOM. **(D)** Boxplot displaying the F-measure for three scRNA-seq “gold/silver standard” data sets. **(E)** Runtime comparisons between PhenoGraph and Seurat. (A & C) Box-and-whisker plots. Boxes represent median ± interquartile range and whiskers ±1.5 × interquartile range. Outliers are represented by black dots.

To test for cell assignment accuracy, we benchmarked FastPG against PhenoGraph, PARC^4^, and FlowSOM^2^. We compared FastPG to PhenoGraph to ensure our method was getting similar accuracy, and FastPG to PARC and FlowSOM due to the published fast speed of these algorithms. Briefly, PARC’s first step employs HNSW, the same kNN approximation as FastPG. PARC next uses a graph-pruning method followed by Leiden^9^ for community detection. FlowSOM applies a self-organizing map for clustering and has faster runtimes due to subsampling.

We evaluated the performance of FastPG using four publicly available “gold standard” mass cytometry data sets from Weber *et al.*^10^; all of which were derived from bone marrow: Levine32 (104,184 cells, 32 protein markers and 14 defined cell types), Levine13 (81,747 cells, 13 protein markers, and 24 defined cell types), Samusik01 (53,173 cells, 37 protein markers, and 24 defined cell types), and SamusikAll (514,386 cells, 37 protein markers, and 24 defined cell types). To better understand the precision and accuracy of FastPG compared to the other methods, we randomly subsampled 20,000 cells from each dataset 20 times. We found that overall, FastPG identified highly similar populations to PhenoGraph compared to the “gold standard” labels as evaluated by the F-measure^2^ (geometric mean of precision and recall, Figure 1B), and both FastPG and PhenoGraph had higher cell assignment accuracy compared to PARC and FlowSOM (median F-measures respectively, Levine13: 0.89, 0.89, 0.88, 0.83; Levine32: 0.79, 0.78, 0.74, 0.79; Samusik01: 0.95, 0.95, 0.87, 0.89; and SamusikAll: 0.95, 0.95, 0.87, 0.87).

We next performed runtime comparisons on a single node containing 64 virtual central processing units (vCPUs; Intel Xeon Skylake CPUs, 2.7 GHz) and 416 GB memory. We randomly sampled cells from a large publicly available dataset (FR-FCM-ZYT6) with a total of 38 million cells to compare the performance of FastPG, PhenoGraph, PARC, and FlowSOM on cell population sizes ranging from 1,000 to 20 million cells. FastPG and FlowSOM were much faster than PhenoGraph and PARC (Figure 1C, 5M cells: 18, 10, 775, 78 minutes, respectively). FastPG was on average 27x faster than PhenoGraph and 7x faster than PARC. FastPG clustered 20 million cells in 1.7 hours, while PARC completed the clustering in 8.6 hours.

The components of PhenoGraph are also implemented as part of Seurat^11^, a popular tool for single cell RNA-seq (scRNA-seq) analysis. To test if FastPG can be applied to cluster scRNA-seq data and provide similar results as Seurat, we tested both methods on three datasets using their “gold/silver standard” cell labels (described in detail in the methods): Zhengmix4eq (4 known cell types, 3,994 cells), Zhengmix8eq (8 known cell types, 3,993 cells), and pbmc68k (11 known cell types, 68,545). Relative to Seurat, we found that FastPG generated almost identical cell assignment accuracy (Figure 1D, median F-measures Seurat and FastPG respectively, Zhengmix4eq: 0.93, 0.92; Zhengmix8eq: 0.76, 0.75; PBMC68k: 0.59, 0.58). FastPG demonstrated greatly increased performance (Figure 1E). FastPG clustered 68,000 cells in 4.3 seconds, whereas Seurat’s runtime was 246.8 seconds. This indicates that while FastPG was developed for mass cytometry data it can be applied to single-cell data in general for quick and accurate determination of cell types.

Together, these experiments suggest that our modifications to the individual components of the PhenoGraph algorithm collectively aid in significantly decreased runtimes with no information loss. FastPG had similar runtime to FlowSOM and higher cell assignment accuracy in three of the four datasets. While PARC had better performance than PhenoGraph, FastPG was much faster and had higher cell assignment accuracy compared to PARC in all four datasets. FastPG allows for fast clustering single cells from population sizes that are previously unmatched for both mass cytometry and scRNA-seq data. This allows for the ability to perform extensive computational experiments for cell assignments, such as performing consensus clustering, and to aid in integration of cells across multiple samples without subsampling.

## Methods

### PhenoGraph and FastPG description

We modified an R implementation of PhenoGraph from Cytofkit^5,12^. PhenoGraph is composed of three main steps: (1) creating a k-nearest neighbor network (kNN) of single cells based using a distance metric calculated from their protein markers, (2) adding weights to the network through the calculation of Jaccard index, and (3) partitioning cells into coherent cell-populations using the Louvain algorithm^6^. Cytofkit uses the space-partitioning kNN method, *k*-dimensional tree, which degrades to linear search with large dimensions^7^. Instead, we used Hierarchical Navigable Small World (HNSW) which has logarithmic scaling due to the hierarchical structure of the search space^7^. We next modified the Jaccard index step to run in parallel using C++ (Rcpp package). Lastly, we replaced the R iGraph implementation of Louvain algorithm with a parallelized version of the Louvain algorithm, Grappolo^8^. Grappolo is a shared-memory parallel software library for community detection implemented in C++ using OpenMP programming model for multithreading. Grappolo implements several parallelization heuristics for fast community detection using the Louvain method as the serial template.

### Comparison of FastPG and PhenoGraph using mass cytometry data

We used FlowCore R package to read in the FCS files using the read.FCS default command.To test cell assignment accuracy of FastPG and PhenoGraph, we set k=30 for the k-Nearest Neighbor step, as was used in Levine *et al.*^5^. We used FastPG v0.0.6 and code from Rphenograph from default settings of Cytofkit last modified on Nov 9, 2017 (https://github.com/JinmiaoChenLab/cytofkit/blob/master/R/Rphenograph.R). We subsampled 20,000 cells without replacement from each of the “gold standard” data sets and used the F measure implementation from the FlowSOM R package. We used the PARC v0.33 using Leidenalg 0.7.0 due to current performance issues with Leidenalg 0.8.0 and FlowSOM v1.18 using default settings, except using k=30 due to the dimensions of the data sets.

For the runtime tests we used a Google Compute Platform instance n1-highmem-64 (64 vCPUs, 416 GB memory). We sampled cells from the large publicly available mass cytometry data set (FR-FCM-ZYT6).

### Comparison of FastPG to PhenoGraph using scRNA-seq data

We used 3 public Unique Molecular Identifier (UMI) count datasets to benchmark FastPG against Seurat. The first two datasets were generated by Duo et al. (2018)^13^; in which florescence-activated cell sorting (FACS)-purified peripheral blood mononuclear cells (PBMCs) from 10X Genomics were equally mixed into four (Zhengmix4eq) and eight (Zhengmix8eq) clusters, respectively. These two datasets serve as gold-standard datasets as the identities of the cells were assigned using FACS, independent from gene expression data. The third dataset, referred to as PBMC68k, contained a large number of PBMCs from one donor^14^. The cell labels for this dataset was based on the gene expression correlation with 11 purified bulk samples. These were used as the “silver standard”

We compared FastPG against Seurat v3.0^11^. For each scRNA-seq dataset, we performed a standardized quality control process using Seurat, with most default parameter settings. We first filtered out genes detected in less than one cell and further excluded cells that had less than 200 detected genes (indication of low-quality) or above 2,500 (indication of doublets). To control for the cell library size effect, normalization was performed on the filtered matrix to obtain the normalized counts. The top 2,000 genes were selected. To reduce the dimensions, principal component analysis was run on the matrix that contained the top 2,000 highly variable genes. Then, we chose the first 30 principal components and k=30 for the kNN for clustering with Seurat and FastPG. We used the Seurat resolution parameter 0.8 to obtain the final cluster labels. We ran FastPG and Seurat subsampling 2,000 cells from the original processed data 20 times for each data set. For clustering performance comparison, we calculated F-measure between the cluster labels and the reference labels implemented from the FlowSOM R package. Time tests were performed on the same GCP instance as the mass cytometry data described above.

## Data Availability

Mass cytometry: Gold standard data sets are available here: https://flowrepository.org/id/FR-FCM-ZZPH. The data set used for testing the runtime performance is available here: https://flowrepository.org/id/FR-FCM-ZYT6 Single-cell RNA-seq: The Duo *et al.* datasets were obtained from the Bioconductor package DuoClustering2018. The Zheng *et al.* dataset was downloaded from https://support.10xgenomics.com/single-cell-gene-expression/datasets.

## Code Availability

FastPG is available here: https://github.com/sararselitsky/FastPG.

All of the code for the analyses are available here: https://github.com/sararselitsky/FastPG_accuracy_performance_scripts

## Acknowledgements

This research was in part supported by the U.S. DOE by the U.S. DOE ExaGraph project, a collaborative effort of U.S. DOE SC and NNSA at Pacific Northwest National Laboratory (PNNL). PNNL is operated by Battelle Memorial Institute under Contract DE-AC06-76RL01830. SRS, JSP, SJ, and TB were supported by Cancer Center Support Grant (P30CA016086) and the University Cancer Research Fund. NS was supported by NIH T32 GM 089626.

